# Brilacidin, a COVID-19 Drug Candidate, demonstrates broad-spectrum antiviral activity against human coronaviruses OC43, 229E and NL63 through targeting both the virus and the host cell

**DOI:** 10.1101/2021.11.04.467344

**Authors:** Yanmei Hu, Hyunil Jo, William F. DeGrado, Jun Wang

## Abstract

Brilacidin, a mimetic of host defense peptides (HDPs), is currently in phase 2 clinical trial as an antibiotic drug candidate. A recent study reported that brilacidin has antiviral activity against SARS-CoV-2 by inactivating the virus. In this work, we discovered an additional mechanism of action of brilacidin by targeting heparan sulfate proteoglycans (HSPGs) on host cell surface. Brilacidin, but not acetyl brilacidin, inhibits the entry of SARS-CoV-2 pseudovirus into multiple cell lines, and heparin, a HSPG mimetic, abolishes the inhibitory activity of brilacidin on SARS-CoV-2 pseudovirus cell entry. In addition, we found that brilacidin has broad-spectrum antiviral activity against multiple human coronaviruses (HCoVs) including HCoV-229E, HCoV-OC43, and HCoV-NL63. Mechanistic studies revealed that brilacidin has a dual antiviral mechanism of action including virucidal activity and binding to coronavirus attachment factor HSPGs on host cell surface. Brilacidin partially loses its antiviral activity when heparin was included in the cell cultures, supporting the host-targeting mechanism. Drug combination therapy showed that brilacidin has a strong synergistic effect with remdesivir against HCoV-OC43 in cell culture. Taken together, this study provides appealing findings for the translational potential of brilacidin as a broad-spectrum antiviral for coronaviruses including SARS-CoV-2.

## INTRODUCTION

Seven coronaviruses are known to infect human beings. Human coronavirus (HCoV)-229E, -NL63, -OC43 and -HKU1 account for 15~30% cases of common cold worldwide ^*1*^, while severe acute respiratory syndrome coronavirus (SARS-CoV) ^*2*^, Middle East Respiratory Syndrome (MERS-CoV) ^*3*^, and severe acute respiratory syndrome coronavirus 2 (SARS-CoV-2)-the causative agent of COVID-19 ^*4*^, are three highly pathogenic human coronaviruses that cause acute severe respiratory syndrome. As the third coronavirus that causes severe respiratory disease, SARS-CoV-2 associated COVID19 has led to more than 211 million infections and over 4.4 million deaths worldwide, and more than 37 million infections and over 627 thousand deaths in the U.S. alone as of Aug 21, 2021 ^*5*^. Currently, two mRNA vaccines are authorized for COVID-19: BNT162b2 (Pfizer, Inc., and BioNTech) and mRNA-1273 (ModernaTX, Inc.), and a third single-dose COVID-19 vaccine JNJ-78436735 (Johnson & Johnson) was issued for Emergency Use Authorization. Although vaccine continues to be a mainstay for viral prophylaxis, the efficacy of vaccine might be compromised with emerging variants such as the delta variant ^*6–8*^. For this reason, small molecular antiviral drugs are important complements of vaccines to help combat pandemics.

Host defense peptides (HDPs), also called antimicrobial peptides (AMPs), are typically small peptides (12-50 amino acids) that are expressed in neutrophils and mucosa and serve as the first line of defense against foreign pathogens ^*9*^. HDPs have been extensively explored as antibiotics ^*10*^, antivirals ^*11*^, antifungals ^*12*^ and anticancer agents ^*13*^. Most HDPs share an amphiphilic structure with a positively charged face and a hydrophobic face ^*14*^. It is proposed that HDPs disrupt bacterial cell membranes by interacting with the negatively charged phospholipid headgroups ^*15–17*^. Brilacidin is a small synthetic HDP mimetic ^*18*^, and has potent antibacterial activity against both Gram-positive and Gram-negative bacteria, and is currently in Phase 2 clinical trials (Clinical Trials NCT01211470, NCT020388, and NCT02324335). The antibacterial mechanisms of action of brilacidin include both membrane disruption and immunomodulation ^*19, 20*^. Brilacidin is also in clinical trial (NCT04784897) as a SARS-CoV-2 antiviral drug candidate for hospitalized COVID-19 patients. A recent study showed that brilacidin exhibited potent inhibitory effect on SARS-CoV-2 replication (EC_50_=0.565 μM/CC_50_=241 μM), and the proposed mechanism of action is through disrupting viral integrity, thereby blocking viral entry ^*21*^. However, the effect of brilacidin on host cell and the antiviral activity of brilacidin against other HCoVs have not been investigated.

In this work, we showed that brilacidin inhibits SARS-CoV-2 pseudovirus entry into multiple cell lines. However, acetyl brilacidin had no inhibition on SARS-CoV-2 pseudovirus entry, and heparin, a heparan sulfate proteoglycans (HSPGs) mimetic, diminished the inhibitory activity of brilacidin. This result suggests that brilacidin has an additional mechanism of action by binding to HSPGs on the host cell, thereby blocking viral attachment. HSPGs have been reported as an attachment factor for SARS-CoV-2 ^*22, 23*^. In addition, we have shown that brilacidin has broad-spectrum antiviral activity against multiple HCoVs including HCoV-229E, HCoV-OC43, and HCoV-NL63. The antiviral mechanism against these viruses similarly involves both virucidal effects and binding to HSPGs. Brilacidin partially loses its antiviral activity against HCoV-229E, HCoV-OC43, HCoV-NL63 in the presence of heparin in cell culture. Drug time-of-addition experiments provided additional evidence that brilacidin exerts its antiviral activity at both viral attachment and early entry stage of viral life cycle. Finally, drug combination therapy demonstrated that brilacidin has strong synergistic effect with remdesivir against HCoV-OC43 in cell culture. Overall, brilacidin appears to have appealing translational potential as a broad-spectrum antiviral for coronaviruses including SARS-CoV-2.

## RESULTS AND DISCUSSION

### Brilacidin inhibits SARS-CoV-2 pseudovirus entry in multiple cell lines

To delineate whether brilacidin blocks SARS-CoV-2 viral entry, we generated pseudotyped HIV-1-derived lentiviral particles with SARS-CoV-2 spike protein ^*24*^, which is widely used to study spike-mediated viral entry into host cells in biosafety level 2 facilities ^*25, 26*^. Brilacidin was tested in SARS-CoV-2 pseudovirus entry assay in several ACE2-expressing cell lines including Vero C1008, Calu-3, Huh-7, Caco-2, and 293T-ACE2. Vero C1008 and 293T-ACE2 express minimal levels of transmembrane serine proteinase 2 (TMPRSS2), therefore SARS-CoV-2 virus enters into these cell lines mainly through endocytosis and relies on endosomal cathepsin L for viral spike protein activation ^*27, 28*^. In contrast, Calu-3 and Caco-2 endogenously express TMPRSS2 ^*29*^, which activates SARS-CoV-2 spike protein on cell surface so virus gets into these cell lines through direct cell membrane fusion. Cathepsin L inhibitor E-64d and TMPRSS2 inhibitor camostat mesylate were included as controls ^*30*^. Our results showed that brilacidin inhibited SARS-CoV-2 pseudovirus entry into all cell lines tested with IC_50_ values ranging from 12.0 ± 1.7 to 23.0 ± 1.6 μM (**Figure 1**). Cytotoxicity assays showed that brilacidin was not toxic to all the cell lines tested at the concentrations examined (**Figure 1F**). Overall, brilacidin inhibits SARS-CoV-2 pseudovirus entry into multiple cell lines. These results suggest the antiviral activity of brilacidin is independent of cathepsin L or TMPRSS2 inhibition.

**Figure 1.**
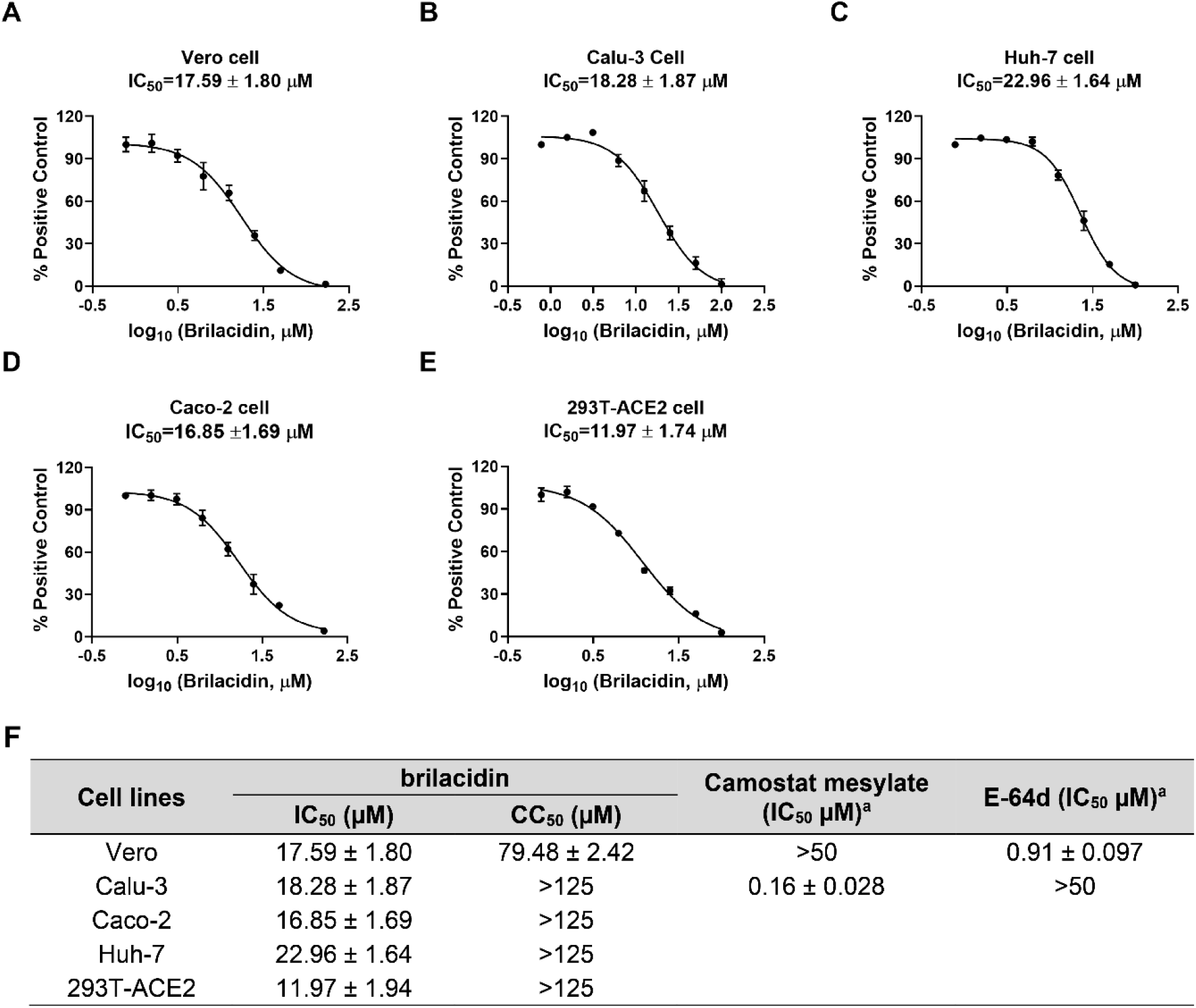
Inhibition of brilacidin on SARS-CoV-2 pseudovirus entry into multiple cell lines including (A) Vero C1008 cells; (B) Calu-3 cells; (C) Huh-7 cells; (D) Caco-2 cells; (E) 293T-ACE2 cells. (F) Cytotoxicity (CC_50_) and inhibitory activity (IC_50_) of brilacidin in SARS-CoV-2 pseudovirus entry assay in different cell lines. ^a^Data from ref ^*31*^. IC_50_ and CC_50_ values were determined through curve fitting described in the “material and methods” section, and all data are mean◻±◻standard deviation of three replicates.

### Brilacidin has broad-spectrum antiviral activity against multiple human coronaviruses, but not influenza or enterovirus

It was recently reported that brilacidin exhibited potent antiviral activity on SARS-CoV-2 replication in both Vero and Calu-3 cells ^*21*^. To test whether brilacidin inhibits the replication of other human coronaviruses, we first tested the antiviral activity of brilacidin against HCoV-229E, HCoV-OC43 and HCoV-NL63 in a viral yield reduction (VYR) assay. The results showed that brilacidin inhibited the replication of HCoV-NL63, HCoV-OC43 and HCoV-229E with EC_50_ values of 2.45 ± 0.05 μM, 4.81 ± 0.95 μM, 1.59 ± 0.07 μM, respectively (**Figure 2A, 2B, 2C**). To confirm the antiviral activity of brilacidin, we tested its inhibitory effect on viral replication over different time course up to 5 days post infection (dpi). HCoV-229E, HCoV-NL63, and HCoV-OC43 were propagated in the presence or absence of brilacidin, and the cell culture supernatants were collected at different time points post infection. Viral titer from each sample was determined by plaque assay. The results demonstrated that brilacidin decreased the viral tiers of all three human coronaviruses by at least 1 log_10_ unit at all time points (**Figure 2D, 2E, 2F**). Brilacidin also inhibited HCoV-OC43 in the plaque assay with an EC_50_ of 7.32 ± 0.15 μM (**Figure 2G, top panel**). In contrast, brilacidin had no effect on the replication of either the influenza A/California/07/2009 (H1N1) virus (**Figure 2G, middle panel**) or enterovirus D68 US/MO/14-18947 (**Figure 2G, bottom panel**) in the plaque assay. The selectivity indices (SI) of brilacidin, which were calculated as the ratio of CC_50_ over EC_50_, range from 17.1 to greater than 64.1 for HCoV-OC43, HCoV-NL63, and HCoV-229E, respectively (**Figure 2H**). Taken together, these results indicate that brilacidin has potent antiviral activity against human coronaviruses, but not influenza or enterovirus D68.

**Figure 2.**
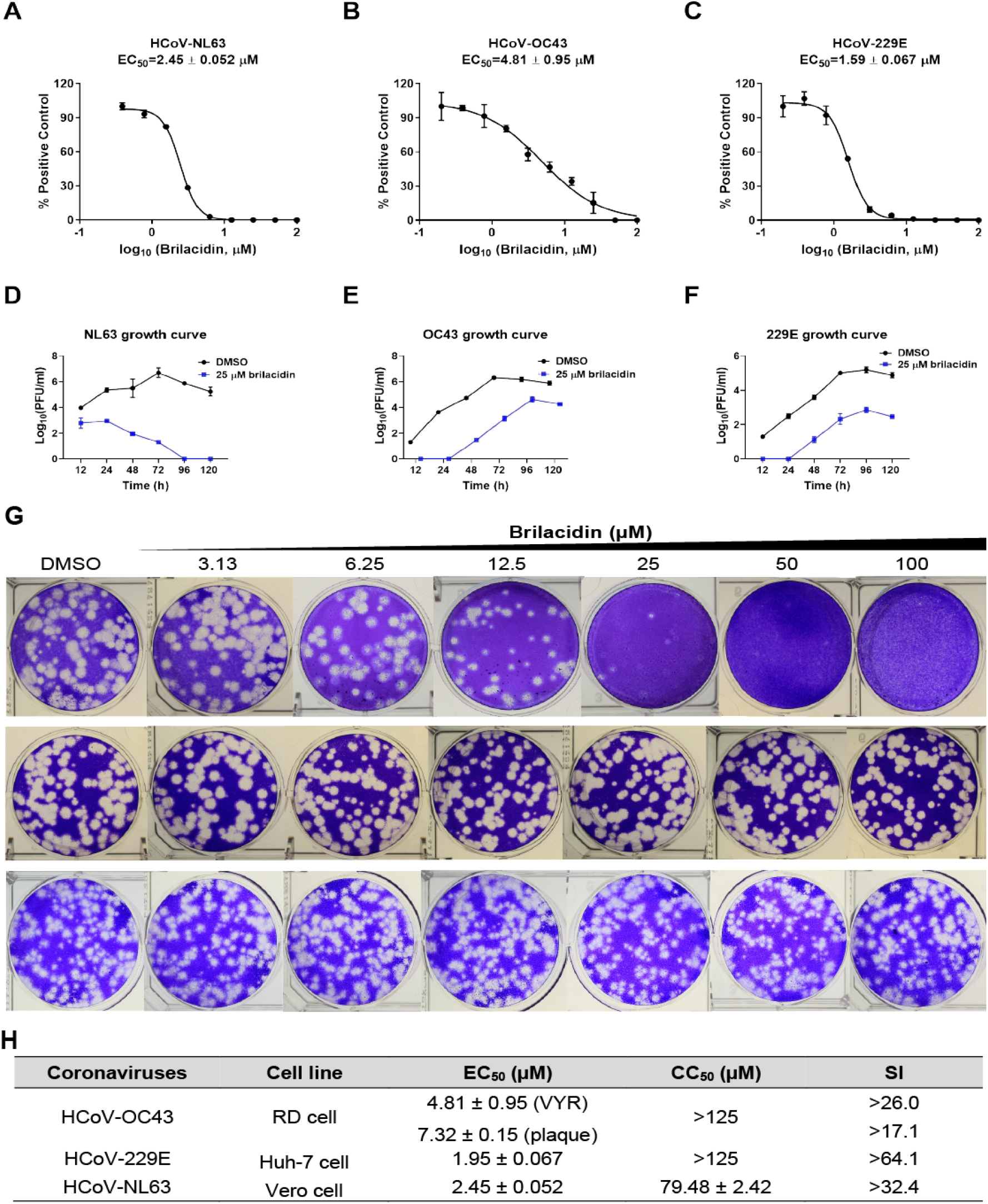
Antiviral activity of brilacidin against multiple human coronaviruses, influenza and enterovirus. Antiviral activity of brilacidin in viral yield reduction assay against HCoV-NL63 (A), HCoV-OC43 (B), and HCoV-229E (C). Growth curve of HCoV-NL63 (D), HCoV-OC43 (E), and HCoV-229E (F) in the presence of DMSO or 25 μM brilacidin. (G) Antiviral activity of brilacidin against HCoV-OC43 (top panel), influenza A/California/07/2009 (H1N1) (middle panel), and enterovirus D68 MO-18947 (bottom panel) in plaque assay. (H) EC_50_ of brilacidin against HCoV-NL63, HCoV-OC43, HCoV-229E; CC_50_ of brilacidin in RD cell, Huh-7 cell, Vero cell; and corresponding SI values. EC_50_ and CC_50_ values were determined through curve fitting described in the “material and methods” section, and all data are mean◻±◻standard deviation of three replicates.

### Brilacidin targets both the virus and the host cell

To elucidate the antiviral mechanism of brilacidin, we first performed experiments to determine whether brilacidin directly targets the virus or the host cell. To assess the virucidal effect of brilacidin on human coronaviruses, we incubated HCoV-OC43, HCoV-229E or HCoV-NL63 with serial concentrations of brilacidin (25, 50, 100, 200 μM) or DMSO at 37 °C for 14 hrs. The mixture was then diluted 10^6^-fold to quantify the infectious viral titer. The final concentrations of brilacidin in each sample after dilution were 0.025, 0.05, 0.1 and 0.2 nM, respectively, which are far below its minimum inhibitory concentration (EC_50_s in the low μM range) and therefore had no effect on plaque formation. It was found that brilacidin treatment decreased the viral titers of all three human coronaviruses dose-dependently, and the viral titers were decreased by more than 1 log_10_ unit at 200 μM (**Figure 3A, 3B, 3C**), meaning over 90% of the viral particles were inactivated by brilacidin treatment. To evaluate the effect of brilacidin on host cells, Huh-7, Vero C1008, and RD cells were pre-treated with serial concentrations of brilacidin (25, 50, 100 μM) or DMSO at 37 °C for 14 hrs, and the cells were subsequently washed with PBS buffer supplemented with magnesium and calcium three times to remove brilacidin. Then the pre-treated cells were infected with HCoV-229E, HCoV-NL63 and HCoV-OC43 at a MOI of 0.1 in the absence of brilacidin. Cell culture supernatants were collected 24 hpi and the viral titers were determined by plaque assay (**Figure 3D, 3E, 3F**). The results demonstrated that pre-treatment of host cells with brilacidin dose-dependently inhibited virus replication and this inhibitory effect is not cell type dependent. Taken together, these results suggest that the antiviral effect of brilacidin involves targeting both the virus and the host cell.

**Figure 3.**
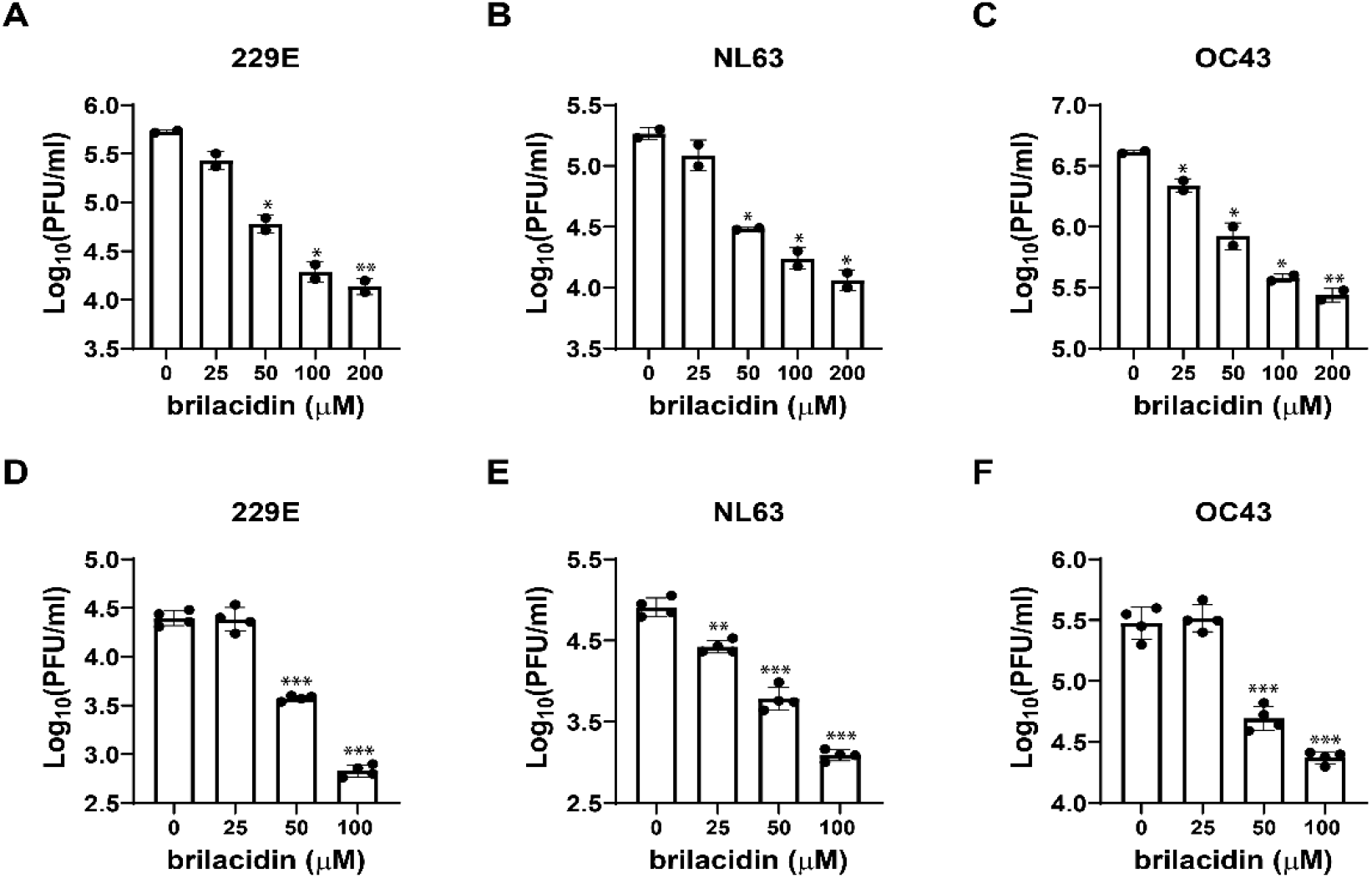
Effect of brilacidin on human coronavirus particles and host cells. Virucidal effect of brilacidin on HCoV-229E (A), HCoV-NL63 (B), and HCoV-OC43 (C). Effects of pre-treatment of cells with brilacidin on the viral replication of HCoV-229E (D), HCoV-NL63 (E), and HCoV-OC43 (F). HCoV-229E, HCoV-NL63, and HCoV-OC43 were propagated in Huh-7, Vero C1008, and RD cells, respectively. *, *p*◻<◻0.05; **, *p*◻<◻0.01; ***, *p*◻<◻0.001 (student’s *t*-test). Data in (A), (B) and (C) are mean◻±◻standard deviation of duplicates, and Data in (D), (E) and (F) are mean◻±◻standard deviation of quadruplicates.

### Brilacidin blocks virus attachment and early entry into host cells

Next, drug time-of-addition experiment was carried out to determine at which step(s) of viral life cycle brilacidin exerts its antiviral activity. In this experiment, 50 μM of brilacidin was added into the cell culture at different time points of viral replication as illustrated in **Figure 4B**. Brilacidin was included in viral attachment and onwards (#1: - 2→14h), viral attachment and entry (#2: -2→0h), viral attachment only (#3: -2→-1h), viral entry and onwards (#4: -1→14h), viral entry only (#5: -1→0h), and different time points post-viral entry (#6-#11). To detect intracellular viral protein levels, RD cells were infected with HCoV-OC43 at an MOI of 1 and cells were fixed 14 hpi for immunofluorescence staining using HCoV-OC43 specific N protein antibody. The immunofluorescence signal was significantly decreased at two time points when brilacidin was added during viral attachment (#1, #2, #3) and the early entry stage (#4, #6, #7) (**Figure 4A**). To quantify progeny viruses released into the cell culture medium, RD cells and Huh-7 cells were infected with HCoV-OC43 and HCoV-229E at an MOI of 0.1, respectively. Viruses in the cell culture medium were collected and the viral titers were determined by plaque assay. Consistent with the immunofluorescence staining results, both HCoV-OC43 and HCoV-229E viral titers decreased considerably when brilacidin was added during the viral attachment (#1, #2, #3) and the early entry stage (#4, #6, #7) (**Figure 4C, 4D**). As shown by the results from both intracellular viral protein level detected by immunofluorescence staining and virus released into cell culture medium quantified by plaque assay, brilacidin exerted the greatest inhibitory effect when it was present at all time points (#1). In conclusion, the drug time-of-addition experiment suggested that brilacidin blocks both viral attachment and early entry into host cells, supporting that it has a dual antiviral mechanism of action.

**Figure 4.**
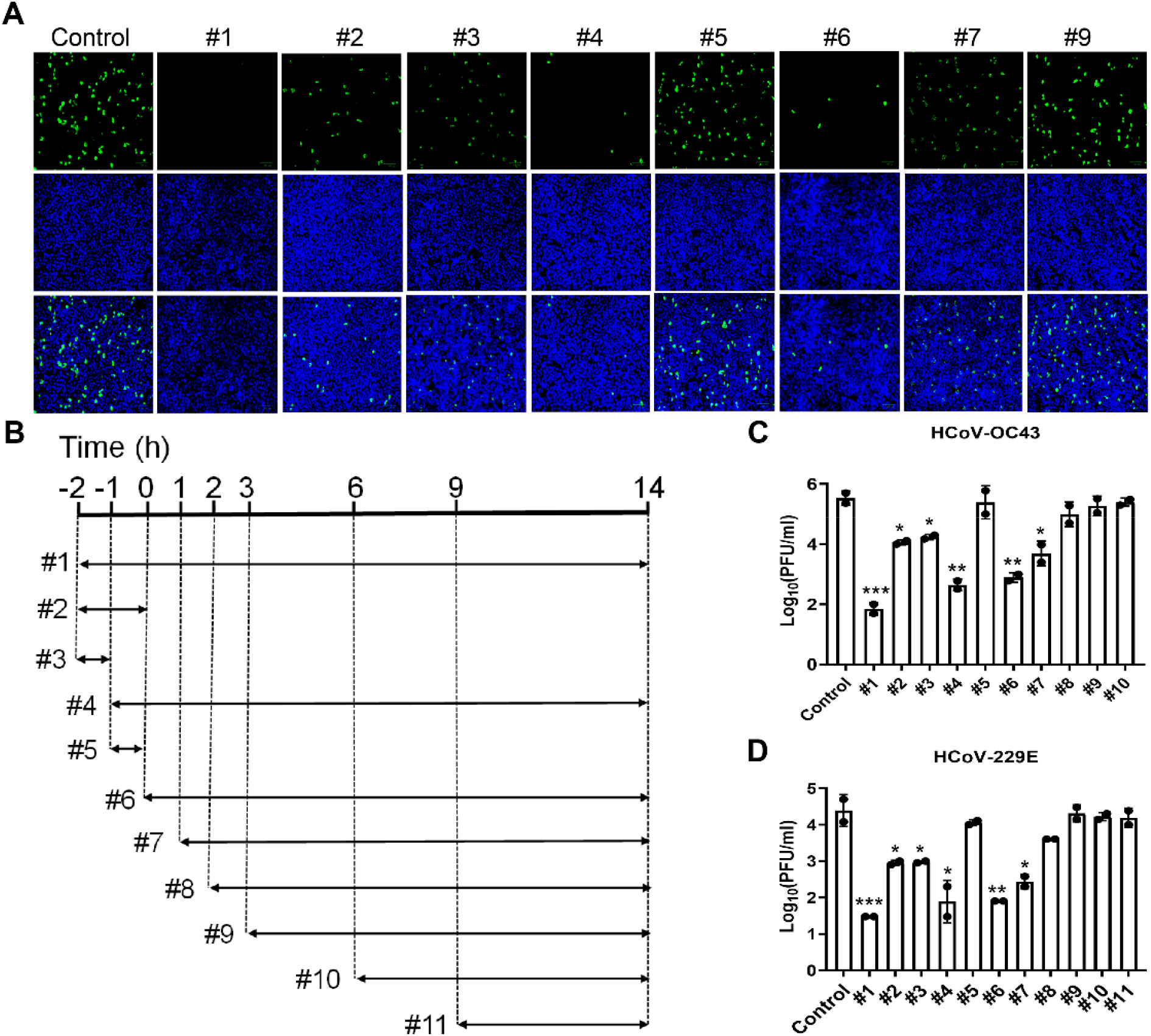
Time-of-addition experiments of brilacidin in inhibiting HCoV-OC43 and HCoV-229E. (A) Representative images of intracellular HCoV-OC43 viral protein detected by immunofluorescence staining using HCoV-OC43 specific antibody. Images were taken by Zoe™ Fluorescent Cell Imager (BioRad); (B) Illustration of the time periods when 50 μM brilacidin was present in the time-of-addition experiments. Arrows represent the periods of time that brilacidin was present in the cell culture; Quantification of HCoV-OC43 (C) or HCoV-229E (D) virus released into the cell culture medium using plaque assay. *, *p*◻<◻0.05; **, *p*◻<◻0.01, ***, *p*◻<◻0.001 (student’s *t*-test). Data are mean◻±◻standard deviation of duplicates.

### Heparin decreases the inhibitory activity of brilacidin in SARS-CoV-2 pseudovirus cell entry and HCoV-OC43, HCoV-229E and HCoV-NL63 replication in cell culture

It was proposed that brilacidin binds to SARS-CoV-2 spike protein ^*30*^. To test whether brilacidin blocks SARS-CoV-2 pseudovirus entry into host cells through interaction with the spike protein, we tested the direct binding of brilacidin to SARS-CoV-2 spike protein receptor binding domain (RBD) using differential scanning fluorimetry (DSF). The results demonstrated that brilacidin has no effect on the melting temperature (*T_m_*) of SARS-CoV-2 spike protein RBD up to 100 μM (**Table 1**), indicating that there is no direct binding between brilacidin and SARS-CoV-2 spike protein RBD.

**Table 1.**
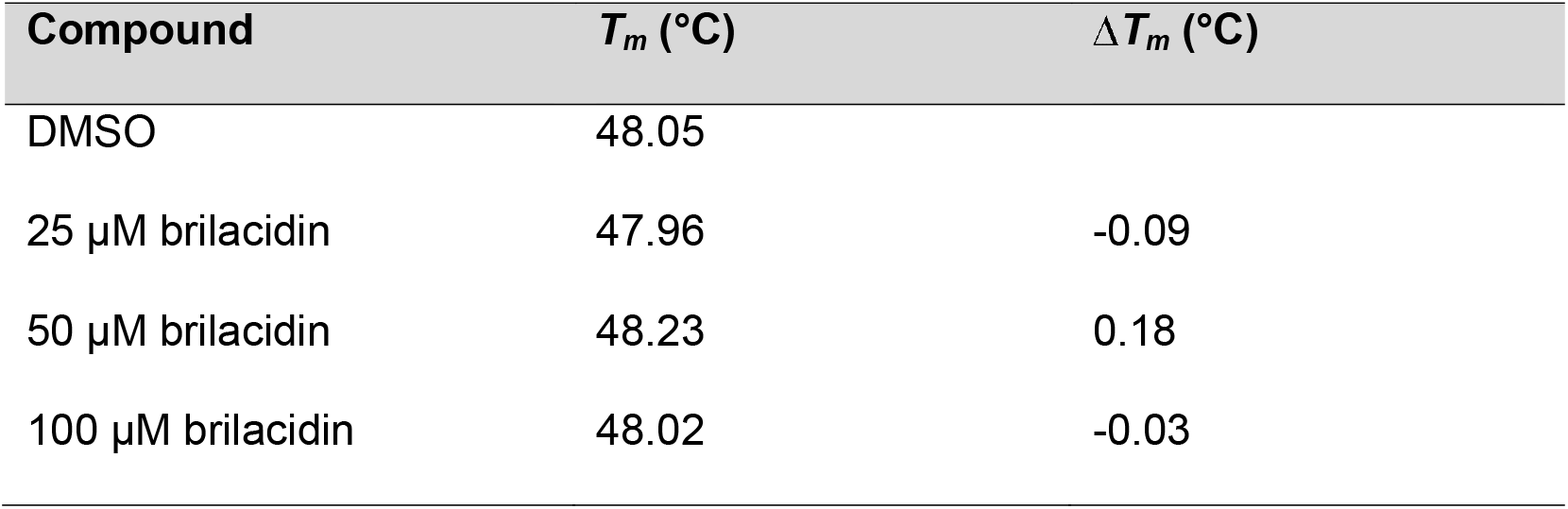
Effect of brilacidin on melting temperature (*T_m_*) of SARS-CoV-2 spike RBD.

| Compound | $T_m$ (°C) | $\Delta T_m$ (°C) |
| --- | --- | --- |
| DMSO | 48.05 |  |
| 25 μM brilacidin | 47.96 | -0.09 |
| 50 μM brilacidin | 48.23 | 0.18 |
| 100 μM brilacidin | 48.02 | -0.03 |

HSPGs are negatively charged, linear polysaccharide that are abundantly expressed on the surface of almost all types of mammalian cells ^*32*^. It has been reported that HCoV-NL63 utilizes cell surface HSPGs as adhesion receptor for viral attachment to target cells through its interaction with the membrane (M) protein ^*33, 34*^. Also, cell surface HSPGs were discovered as the attachment factors for SARS-CoV-2 and facilitate the subsequent binding of spike protein to ACE2 receptor ^*35–37*^. Brilacidin is +4 charged at neutral pH, we therefore hypothesize that brilacidin might bind to the cell surface HSPGs through electrostatic interactions, thereby blocking viral attachment and entry. To test this hypothesis, we chose acetyl brilacidin, which is +2 charged, as a control compound (**Figure 5A**). It was found that acetyl brilacidin completely lost inhibitory activity in SARS-CoV-2 pseudovirus entry assay in Vero C1008, Calu-3 and Caco-2 cells (**Figure 5B, 5C, 5D**). This result suggests that the +4 charge on brilacidin is critical for the antiviral activity, and the antiviral mechanism of action might involve interaction with the binding to HSPGs.

**Figure 5.**
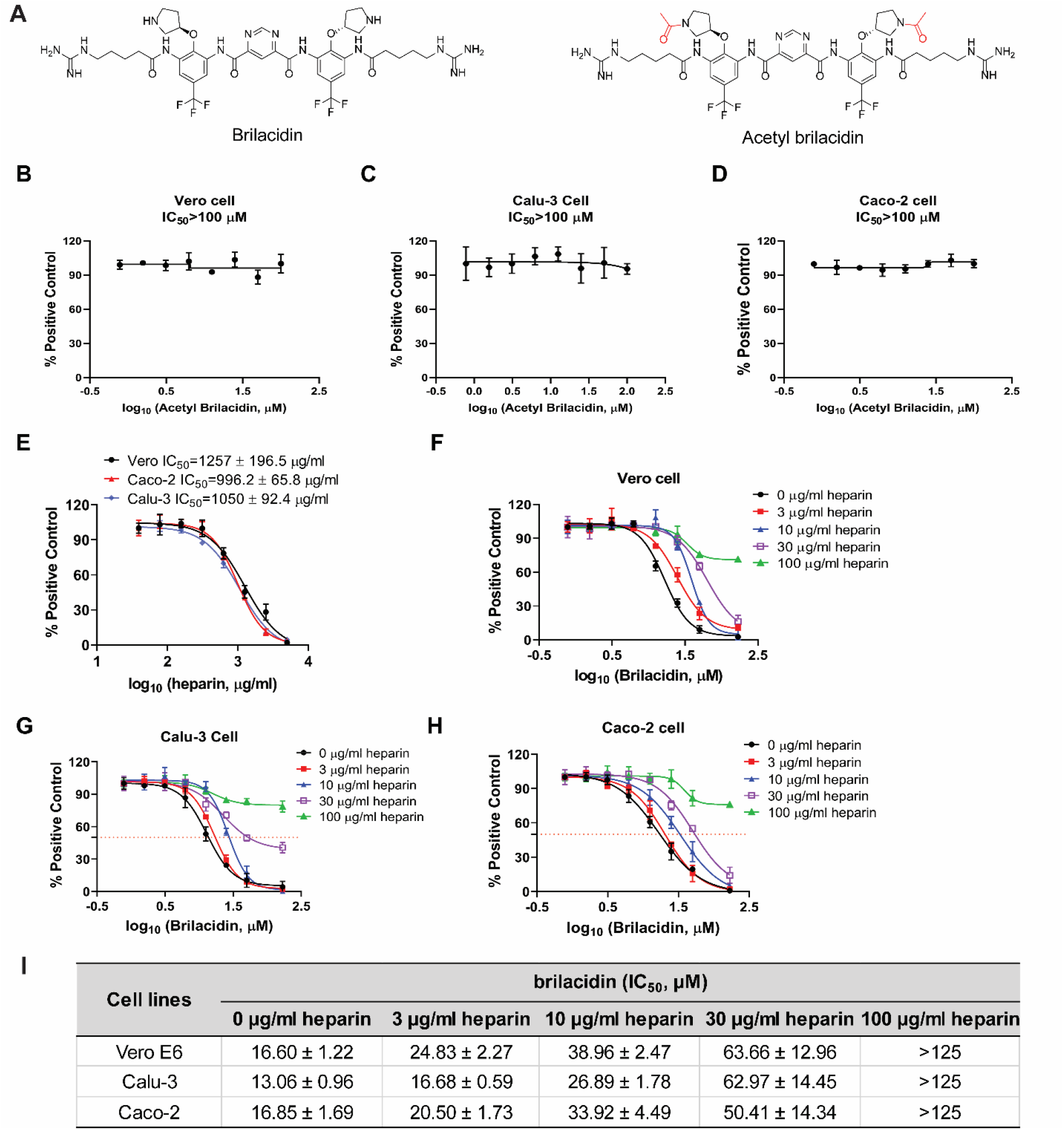
Heparin diminishes the inhibitory activity of brilacidin in SARS-CoV-2 pseudovirus entry into different cell lines. (A) Chemical structures of brilacidin and acetyl brilacidin. Effect of acetyl brilacidin on SARS-CoV-2 pseudovirus entry into Vero C1008 (B), Calu-3 (C), and Caco-2 (D). (E) IC_50_s of heparin in SARS-CoV-2 pseudovirus entry assay in Vero C1008 cells, Caco-2 cells and Calu-3 cells. Heparin dose-dependently decreased the potency of brilacidin in inhibiting SARS-CoV-2 pseudovirus entry into Vero C1008 cells (F), Calu-3 cells (G), and Caco-2 cells (H). (I) Summary of IC_50_s of brilacidin in SARS-CoV-2 pseudovirus entry assay in the heparin competition assay. IC_50_ values were determined through curve fitting described in the “material and methods” section, and all data are mean ± SD of two independent experiments.

If brilacidin binds to cell surface HSPGs in cell culture, exogenous addition of HSPG mimetics such as heparin will compete with HSPGs for binding of brilacidin, resulting in decreased inhibitory activity of brilacidin on SARS-CoV-2 pseudovirus entry and the replication of human coronaviruses in cell culture. We therefore performed the competition assay to evaluate the effect of heparin on the antiviral activity of brilacidin. To test the effect of heparin on brilacidin activity in SARS-CoV-2 pseudovirus entry, heparin was first tested in SARS-CoV-2 pseudovirus entry assay in Vero C1008, Caco-2 and Calu-3 cells to determine proper concentrations in the competition assay. The highest concentration of heparin used in the competition assay was 100 μg/ml, which had no effect on SARS-CoV-2 pseudovirus entry (**Figure 5E**). As expected, addition of heparin dose-dependently abolished the inhibitory activity of brilacidin on SARS-CoV-2 pseudovirus entry into Vero C1008 cells (**Figure 5F**), Calu-3 cells (**Figure 5G**), and Caco-2 cells (**Figure 5H**) as shown by the increasing IC_50_ values. Specifically, heparin increased the IC_50_ values of brilacidin by more than 2-fold and 3 to 5-fold at 10 and 30 μg/ml, respectively (**Figure 5I**). Heparin almost completely abolished the inhibitory activity of brilacidin when added at 100 μg/ml concentration (IC_50_ > 125 μM) (**Figure 5E, 5F, 5G, 5H**).

To test whether heparin affects the inhibition of human coronaviruses by brilacidin, HCoV-229E, HCoV-NL63 and HCoV-OC43 were amplified in the presence of different concentrations of brilacidin alone or combination of brilacidin and heparin. The intracellular viral level of amplified HCoV-OC43 was detected by immunofluorescence staining (**Figure 6A, 6B**), and the amplified HCoV-229E, HCoV-NL63 and HCoV-OC43 viruses released into culture medium were quantified by plaque assay (**Figure 6C, 6D, 6E**). Consistent with previous results, brilacidin dose-dependently inhibited replication of all three HCoVs (#1 vs #5 and #9). Addition of heparin dose-dependently decreased the inhibitory activity of brilacidin on replication of all three HCoVs (#5 vs #7 and #8; #9 vs #11 and #12).

**Figure 6.**
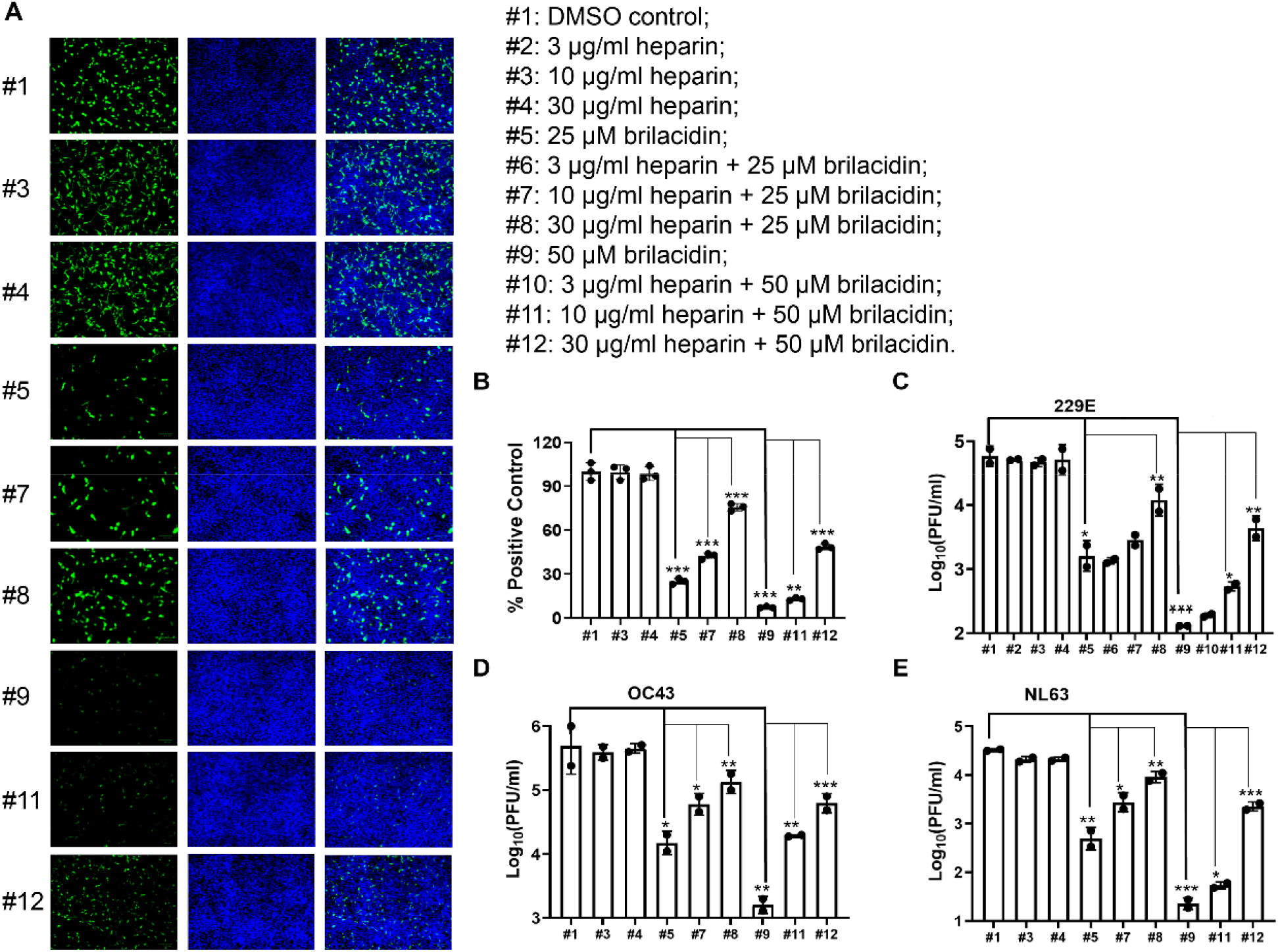
Heparin decreases the inhibitory activity of brilacidin on HCoVs replication in cell culture. RD cell, Huh-7 cell and Vero C1008 cell were infected with HCoV-OC43, HCoV-229E and HCoV-NL63 at an MOI of 0.1, respectively. Cell culture medium were collected 24 hpi to determine viral titers in each sample. RD cells were fixed 24 hpi for immunofluorescence staining, and intracellular HCoV-OC43 viral protein level was detected by HCoV-OC43 specific antibody. (A) Representative images of intracellular HCoV-OC43 viral protein level in RD cells detected by immunofluorescence staining; (B) Quantification of HCoV-OC43 viral protein level from panel A. Three groups (five for each group) of images were captured from three different areas in each sample, and fluorescent signals were quantified in Image J by calculating the percentage of viral protein fluorescent signal (green) to nuclei fluorescent signal (blue) in pixels. Results shown are the average percentages from all three groups and normalized to DMSO control. Viral titers of viruses released into cell culture from each sample of (C) HCoV-OC43; (D) HCoV-229E; (E) HCoV-NL63. *, *p*◻<◻0.05; **, *p*◻<◻0.01; ***, *p*◻<◻0.001 (student’s *t*-test). All data are mean ± SD of two independent experiments.

### Brilacidin has a strong synergistic antiviral effect with remdesivir in cell culture

Combination therapy is commonly used to slow down drug resistance development and reduce side effects ^*38, 39*^. The antiviral effect of brilacidin and remdesivir in combination therapy was evaluated in HCoV-OC43 plaque assay using the combination indices method (**Figure 7**) ^*31*^. Remdesivir, a SARS-CoV-2 polymerase inhibitor, is the only FDA-approved antiviral for treating COVID-19. Brilacidin and remdesivir were mixed at different ratios and the corresponding EC_50_ values for brilacidin and remdesivir were calculated. Combination indices (CIs) versus the EC_50_ values of brilacidin and remdesivir at different combination ratios were plotted (**Figure 7A**). The red line indicates additive effect; the right upper area above the red line indicates antagonism, while the left bottom area below the red line indicates synergy ^*37*^. The CIs at all the combination ratios fell below the red line (**Figure 7A**), and the fractional inhibitory concentration index (**FICI**) which was used to determine synergistic effects of compounds are less than 0.5 at all combination ratios (**Figure 7B**), suggesting brilacidin has significant synergistic antiviral effect with remdesivir in the combination therapy.

**Figure 7.**
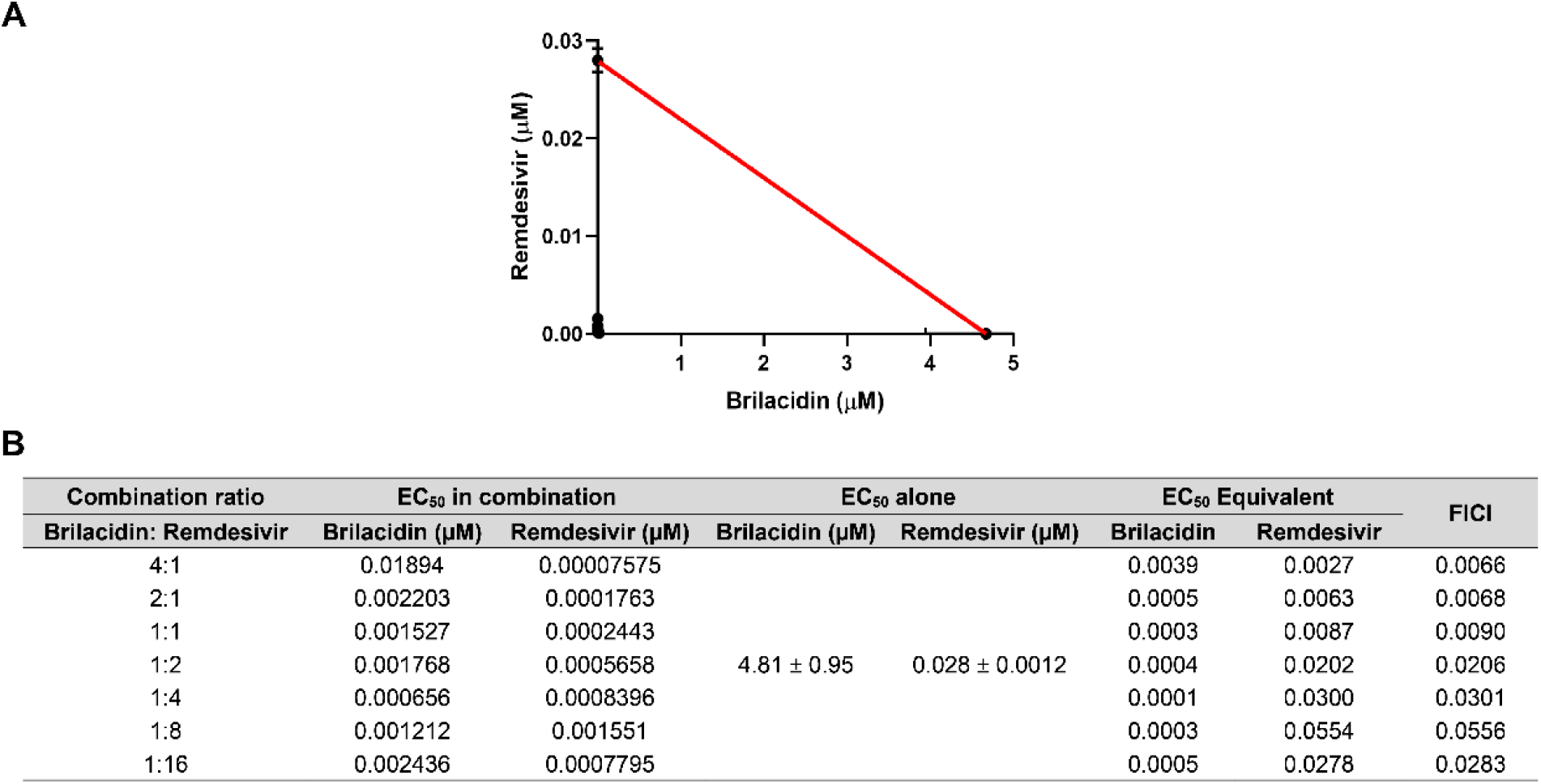
Combination therapy of brilacidin with remdesivir in cell culture. (A) Plot of combination indices (CIs) versus the EC_50_ values of brilacidin and remdesivir at different combination ratios. (B) Table of combination therapy with EC_50_ and FICI values. EC_50_ equivalent was the ratio of EC_50_ of the compound in each combination to its EC_50_ alone. FICI was the sum of brilacidin and remdesivir EC_50_ equivalent in each combination. Data are mean ± SD of two independent experiments.

## CONCLUSION

As the COVID19 pandemic keeps ongoing and variants continue to emerge, effective therapeutic interventions are urgently needed. Although three vaccines are currently available for prevention of COVID19, there is an urgent need for small molecular antivirals to help combat the pandemic. In this study, we investigated the antiviral activity and mechanism of action of brilacidin against multiple human coronaviruses. Our findings include: 1) Brilacidin has broad-spectrum antiviral activity against HCoV-OC43, HCoV-NL63, and HCoV-229E viruses in cell culture; 2) Brilacidin inhibits SARS-CoV-2 pseudovirus entry into multiple cell lines, indicating that the inhibition is not cell type dependent; 3) Brilacidin has dual antiviral mechanisms of action which involves targeting both the virus and the host cell. Brilacidin has virucidal activity and blocks viral attachment to host cells by binding to HSPGs; 4) Brilacidin has strong synergistic antiviral effect with the FDA-approved SARS-CoV-2 antiviral remdesivir against HCoV-OC43 in cell culture.

The proposed antiviral mechanism of brilacidin is summarized in a model illustrated in **Figure 8**, which is supported by multiple lines of evidence. Our results showed that brilacidin has a dual antiviral mechanism against human coronaviruses including blocking viral attachment to host cells through binding to HSPGs and virucidal activity. HCoV-OC43, HCoV-NL63 and HCoV-229E showed dose-dependent decrease of replication in cells pretreated with brilacidin, and viral particles lose infectivity after incubation with brilacidin (**Figure 3**). Drug time-of-addition experiment suggested that brilacidin exerted its antiviral activity at two individual steps: viral attachment to host cell and early entry after entering into the host cells (**Figure 4**). The inhibition of viral attachment by brilacidin was confirmed in SARS-CoV-2 pseudovirus entry assay (**Figure 1**). DSF assay results demonstrated that brilacidin has no direct binding to SARS-CoV-2 spike protein RBD (**Table 1**). Competition experiment with heparin indicated that brilacidin binds to host cell surface HSPGs to block viral attachment to host cells. Addition of heparin dose-dependently decreased the inhibition of brilacidin in SARS-CoV-2 pseudovirus entry assay (**Figure 5**) and the replication of HCoV-OC43, HCoV-NL63 and HCoV-229E in cell culture (**Figure 6**).

**Figure 8.**
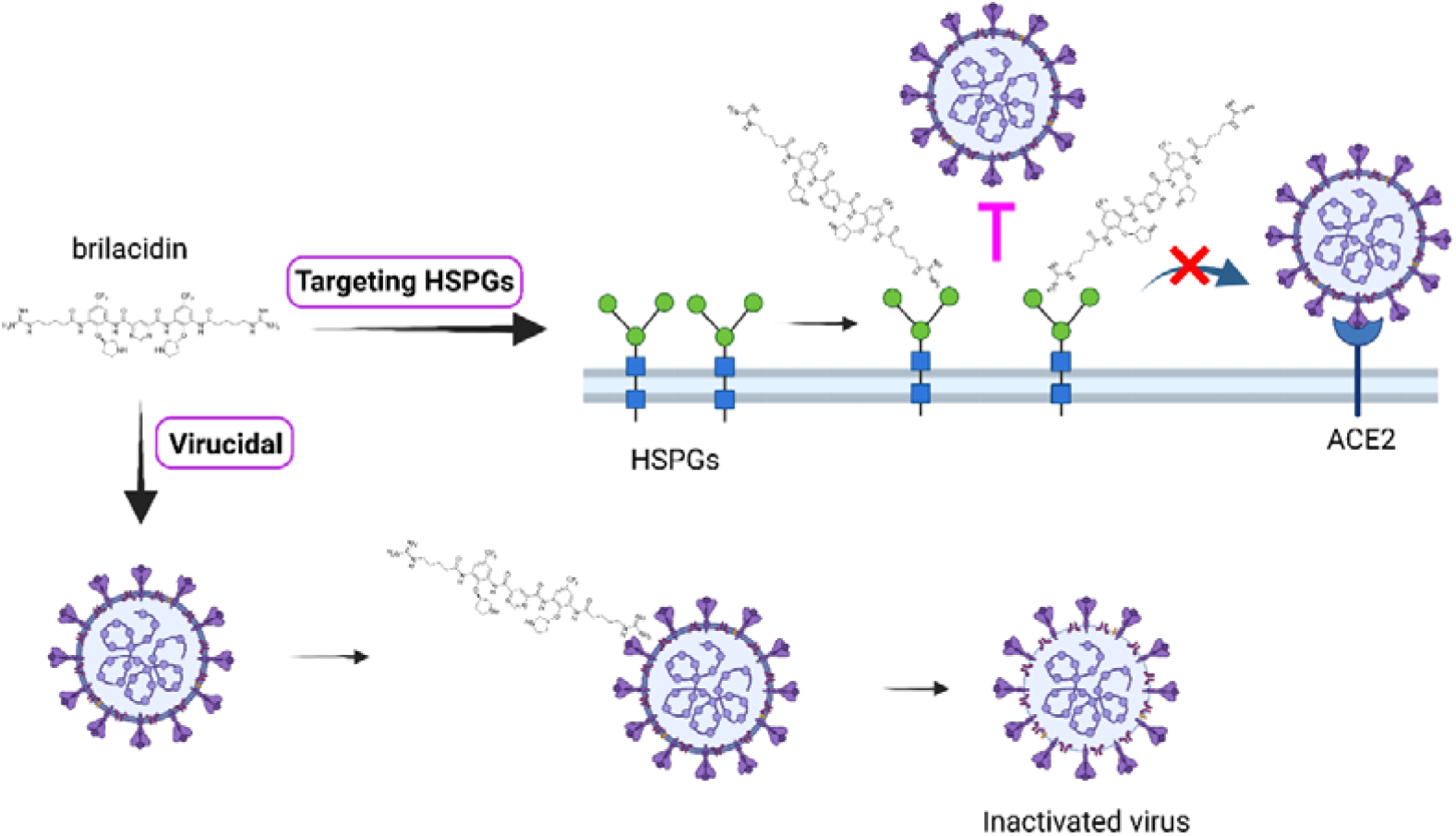
Proposed antiviral mechanism of brilacidin. Brilacidin has a dual antiviral mechanism including block viral attachment to host cells through binding to HSPGs and disrupt viral particles. The figure was created with BioRender.com.

In summary, our results indicate that brilacidin has a dual antiviral mechanism of action including targeting host cell surface HSPGs to block viral attachment and inactivating viral particles. This dual antiviral mechanism of action might slow down the pace of resistance development. Taken together, the broad-spectrum antiviral activity of brilacidin against coronaviruses warrants its further development as a broad-spectrum antiviral for the treatment of not only current COVID-19 but also future emerging coronaviruses.

## METHORDS

### Cell lines, viruses and reagents

Human rhabdomyosarcoma cell line (RD, ATCC^®^ CCL-136™), African green monkey kidney cell line Vero C1008 (ATCC^®^ CRL-1586™), Human hepatoma cell line Huh-7 (a kind gift from Dr. Tianyi Wang at University of Pittsburgh), and HEK293T expressing ACE2 (293T-ACE2, BEI Resources, NR-52511) cell lines were maintained in Dulbecco’s modified eagle’s medium (DMEM); Human fibroblast cell line MRC-5 (ATCC^®^ CCL-171™), Human lung adenocarcinoma cell line Calu-3 (ATCC^®^ HTB-55™), human Colorectal adenocarcinoma cell line (Caco-2, ATCC® HTB-37™) were maintained in eagle’s minimum essential medium (EMEM, ATCC® 30-2003™). Both media were supplemented with 10% fetal bovine serum (FBS) and 1% penicillin-streptomycin antibiotics. Cells were kept at cell culture incubator (humidified, 5% CO_2_/95% air, 37 °C). The following reagents were obtained through BEI Resources, NIAID, NIH: human coronavirus, HCoV-OC43, NR-52725; human coronavirus, HCoV-NL63, NR-470. HCoV-OC43 was propagated in RD cell line; HCoV-NL63 was initially propagated in 293T-ACE2 cell line and accommodated in Vero E6 cell line. HCoV-229E was obtained from Dr. Bart Tarbet (Utah State University) and amplified in Huh-7 or MRC-5 cell lines.

### Antiviral assays

The antiviral activity of brilacidin was tested against HCoV-229E, HCoV-NL63 and HCoV-OC43 in Viral yield reduction (VYR) assays as previously described ^*30, 40–42*^. Briefly, viruses were first replicated in the presence of serial concentrations of brilacidin (0, 0.39, 0.78, 1.56, 3.13, 6.25, 12.5, 25, 50, 100 μM). Progeny virions released in the supernatant were collected 24 hrs post-infection from each concentration of brilacidin and the viral titers were determined by plaque reduction assay. Viruses were serially diluted 10 to 10^6^ folds and infect the cells in 6-well plate. The infected cells were incubated at 33 or 37 °C for 1 h to allow virus adsorption. The viral inoculum was removed and an overlay containing 0.6% Avicel supplemented with 2% FBS in DMEM was added and incubated in the 33 or 37 °C incubator for 4 to 5 days. The plaque formation was detected by staining the cell monolayer with crystal violet. HCoV-229E and HCoV-OC43 plaque assays were carried out on RD cells and incubated at 33 °C, HCoV-NL63 plaque assay was performed on Vero C1008 cells and incubated at 37 °C. EC_50_ values were determined by plotting percentage of positive control versus log_10_ compound concentrations from best-fit dose response curves with variable slope in Prism 8.

Viral growth curves were obtained by replicating viruses in the presence or absence of 25 μM brilacidin at MOI of 0.1. Viruses in the supernatant were collected at the indicated time point post infection and viral titers were determined by plaque reduction assay as described in VYR assay section.

The antiviral activity of brilacidin tested in HCoV-OC43 plaque assay was carried out similarly as described in VYR assay, except that about 100 PFU of HCoV-OC43 virus was used to infect the cells in each well of 6-well plate and serial concentrations of brilacidin (0, 3.13, 6.25, 12.5, 25, 50, 100 μM) was included in the Avicel overlay. The plaque areas were quantified using Image J and the EC_50_ value was determined by plotting percentage of plaque area versus log_10_ compound concentrations from best-fit dose response curves with variable slope in Prism 8.

The antiviral activity of brilacidin against influenza and enterovirus D68 was carried out in plaque assay as previously described ^*36, 43, 44*^.

### Cytotoxicity assay

Cytotoxicity of brilacidin was evaluated in different cell lines using the neutral red uptake assay as previously described ^*45, 46*^. Cells were dispensed into 96-well plate at a density of 1×10^5^ cells/ml at 100 μl/well. The growth medium was removed 18-24 hrs later and the cells were washed with 200 μl PBS supplemented with magnesium and calcium, and 200 μl fresh medium (+2% FBS) containing serial concentrations of brilacidin (0, 1.9, 3.9, 7.8, 15.6, 31.3, 62.5, 125 μM) was added into each well. After incubating at 37 °C incubator with 5% CO_2_ for 48 hrs, cells were stained with 40 μg/ml neutral red for 2-4 hrs at 37 °C. The amount of neutral red uptaken by live cells was quantified by measuring the absorbance at 540 nm using a Multiskan FC Microplate Photometer (Fisher Scientific). The CC_50_ values were determined from best-fit dose response curves with variable slope in Prism 8.

### Time of addition

Drug time-of-addition experiment was performed as previously described ^*30, 47*^. Briefly, RD cells were seeded at 1 ×10^5^ cells/well in 12-well plate with cover slip (Cat#: GG-12-1.5-PDL, Neuvitro) and infected with HCoV-OC43 at MOI of 1 and 25 μM brilacidin was added at different time points of viral life cycle as illustrated in Figure 3B. At 14 h post infection (hpi), cells were fixed with 4% formaldehyde for 10 min followed by permeabilization with 0.2% Triton X-100 for another 10 min. After blocking with 5% bovine serum, cells were sequentially stained with mouse anti-Coronavirus antibody, HCoV-OC43 strain, clone 541-8F (Cat#: MAB9012, Millipore Sigma, Burlington, Massachusetts, USA) as primary antibody, and anti-mouse secondary antibody conjugated to Alexa-488 or Alexa546 (Cat # A-11029, Cat # A-11030, Thermo Scientific, Waltham, Massachusetts, USA). Nuclei were stained with 300 nM DAPI (Cat#: D1306, Thermo Scientific, Waltham, Massachusetts, USA) after secondary antibody incubation. For time of addition experiment using HCoV-OC43 and HCoV-229E in plaque assay, RD cells or Huh-7 cells were infected at MOI of 0.1 and 25 μM brilacidin was added at different time points of viral life cycle. Progeny virions released into the supernatant were harvested at 14 hpi and the viral titers were determined by plaque assay.

### Pseudovirus assay

A pseudotype HIV-1-derived lentiviral particles bearing SARS-CoV-2 Spike and a lentiviral backbone plasmid encoding luciferase as reporter was produced in HEK293 T cells engineered to express the SARS-CoV-2 receptor ACE2 (293 T-ACE2 cells), as previously described ^*24*^. The pseudovirus was then used to infect Vero C1008 cells, Huh-7 cells, Caco-2 cells, Calu-3 cells or 293 T-ACE2 cells in 96-well plates in the presence of serial concentrations of brilacidin (0, 3.13, 6.25, 12.5, 25, 50, 100 μM). Cells were lysed 48 hpi using the Bright-Glo Luciferase Assay System (Cat#: E2610, Promega, Madison, WI, USA), and the cell lysates were transferred to 96-well Costar flat-bottom luminometer plates. The relative luciferase units (RLUs) in each well were detected using Cytation 5 Cell Imaging Multi-Mode Reader (BioTek, Winooski, VT, USA). The IC_50_ values were determined from best-fit dose response curves with variable slope in Prism 8.

### Differential scanning fluorimetry (DSF)

Direct binding of brilacidin with SARS-CoV-2 Spike protein receptor binding domain (RBD) was detected by differential scanning fluorimetry (DSF) using a Thermal Fisher QuantStudio 5 Real-Time PCR System as previously described ^*48, 49*^ with minor modifications. SARS-CoV-2 (2019-nCoV) Spike RBD-His Recombinant Protein (Cat. #: 40592-V08H, SinoBiological) was diluted in PBS buffer to a final concentration of 4 μM, and incubated with serial concentrations of brilacidin (25, 50, 100 μM) at 30 °C for 1 hr. DMSO was included as reference. 1× SYPRO orange (Thermal Fisher, Cat. #: S6650) was added and the fluorescence signal was recorded under a temperature gradient ranging from 20 to 95 °C (incremental step of 0.05 °C s^−1^). The melting temperature (*T_m_*) was calculated as mid log of the transition phase of the protein from the native to the denatured state using a Boltzmann model in Protein Thermal Shift Software v1.3. Δ *T_m_* was calculated by subtracting melting temperature of protein in the presence of DMSO from the melting temperature of proteins in the presence of brilacidin.

### Combination therapy

The combination antiviral effects of brilacidin and remdesivir were evaluated in HCoV-OC43 plaque assay in cell culture. Brilacidin was mixed with remdesivir at fixed EC_50_ ratios of 4:1, 2:1, 1:1, 1:2, 1:4, 1:8, and 1:16 separately. In each combination, nine 3-fold serial dilutions (equal to a 0.5 log_10_ unit decrease) of brilacidin and remdesiivr mixture were tested to plot the dose inhibition curve, based on which the EC_50_ values of individual brilacidin and remdesivir were determined in each combination. A combination indices (CIs) plot was used to depict the EC_50_ values of brilacidin and remdesivir at different combination ratios. The red line indicates the additive effect, and above the red line indicates the antagonism, while below the red line indicates the synergy ^*50*^. The fractional inhibitory concentration index (FICI) was calculated using the following formula: FICI = [(EC_50_ of brilacidin in combination)/(EC_50_ of brilacidin alone)] + [(EC_50_ of remdesivir in combination)/(EC_50_ of remdesivir alone)]. FICI <0.5 was interpreted as a significant synergistic antiviral effect ^*51*^.

## AUTHOR INFORMATION

### Corresponding Author

**Jun Wang** – Department of Pharmacology and Toxicology, College of Pharmacy, The University of Arizona, Tucson, Arizona 85721, United States; Phone: +1-520-626-1366;

### Authors

**Yanmei Hu** – Department of Pharmacology and Toxicology, College of Pharmacy, The University of Arizona, Tucson, Arizona 85721, United States

**Hyunil Jo** - Department of Pharmaceutical Chemistry, School of Pharmacy, University of California, San Francisco, California 94158, United States

**William F. DeGrado** - Department of Pharmaceutical Chemistry, School of Pharmacy, University of California, San Francisco, California 94158, United States

## Author Contributions

J.W. and W.F.D. conceived and designed the study; Y.H. performed the pseudovirus neutralization assay, antiviral assays, time of addition experiment, immunofluorescence assays, thermal shift binding assay, and the combination therapy experiment. H. J. provided the brilacidin and acetyl brilacidin samples. Y.H. and J.W. wrote the manuscript.

## Notes

The authors declare no competing financial interest(s).

## ACKNOWLEDGMENTS

This research was partially supported by the National Institute of Allergy and Infectious Diseasess of Health (NIH) (grants AI147325, AI157046, and AI158775) and the Arizona Biomedical Research Commission Centre Young Investigator grant (ADHS18-198859) to J. W. Y.H. was supported by the NIH training grant T32 GM008804.

---

For Table of Contents Use Only

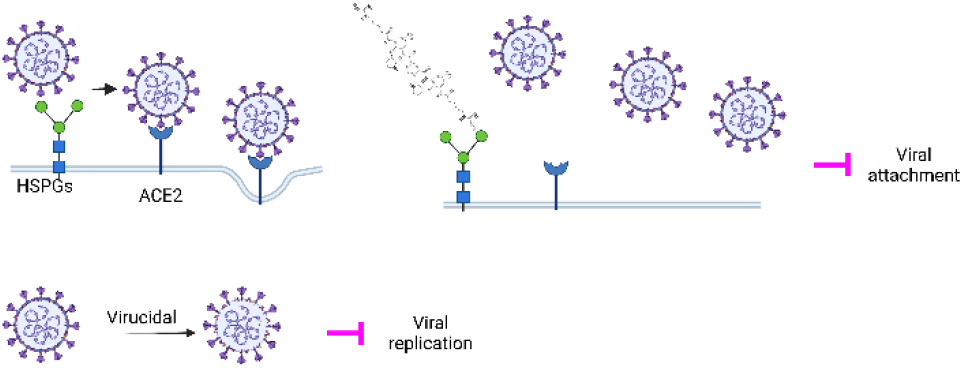
Brilacidin inhibits human coronaviruses through blocking viral attachment to HSPGs and inactivating the virus.

## Notes

### Competing Interest Statement

The authors have declared no competing interest.

